# SVEngine: an efficient and versatile simulator of genome structural variations with features of cancer clonal evolution

**DOI:** 10.1101/247536

**Authors:** Li Charlie Xia, Dongmei Ai, Hojoon Lee, Noemi Andor, Chao Li, Nancy R. Zhang, Hanlee P. Ji

## Abstract

**Background:** Simulating genome sequence data with features can facilitate the development and benchmarking of structural variant analysis programs. However, there are a limited number of data simulators that provide structural variants in silico. Moreover, there are a paucity of programs that generate structural variants with different allelic fraction and haplotypes.

**Findings:** We developed SVEngine, an open source tool to address this need. SVEngine simulates next generation sequencing data with embedded structural variations. As input, SVEngine takes template haploid sequences (FASTA) and an external variant file, a variant distribution file and/or a clonal phylogeny tree file (NEWICK) as input. Subsequently, it simulates and outputs sequence contigs (FASTAs), sequence reads (FASTQs) and/or post-alignment files (BAMs). All of the files contain the desired variants, along with BED files containing the ground truth. SVEngine’s flexible design process enables one to specify size, position, and allelic fraction for deletion, insertion, duplication, inversion and translocation variants. Finally, SVEngine simulates sequence data that replicates the characteristics of a sequencing library with mixed sizes of DNA insert molecules. To improve the compute speed, SVEngine is highly parallelized to reduce the simulation time.

**Conclusions:** We demonstrated the versatile features of SVEngine and its improved runtime comparisons with other available simulators. SVEngine’s features include the simulation of locus-specific variant frequency designed to mimic the phylogeny of cancer clonal evolution. We validated the accuracy of the simulations. Our evaluation included checking various sequencing mapping features such as coverage change, read clipping, insert size shift and neighbouring hanging read pairs for representative variant types. SVEngine is implemented as a standard Python package and is freely available for academic use at: https://bitbucket.org/charade/svengine.

## FINDINGS

### Background

Next generation sequencing (**NGS**) has enabled researchers to detect and resolve complex genomic structural features at base-pair resolution. One can detect a variety of structural variations **(SVs)** including deletions, insertions, inversions, tandem duplications and translocations based on NGS whole genome sequence data [1]. A variety of algorithms have been developed for structural variant calling from NGS data. This includes programs such as Breakdancer, CNVnator, Delly, Haplotype Caller, Lumpy, SWAN, Pindel among others [2–10]. Even with these programs, accurate SV detection remains a significant challenge. For example, some SVs occur in lower allelic fractions as seen in tumors with intratumoral heterogeneity [11]. This is frequently the case in sequencing tumor samples, where cancer starts from a seeding clone and through clonal evolution, successively acquires additional rearrangements at lower allelic fractions.

Benchmarking structural variant callers with available ground truth data sets is critical for software tool development, bioinformatics pipeline testing and objective assessment of detection accuracy [12]. Whole genome data sets are available from high sequencing coverage with Illumina or Pacific Bioscience systems [13]. However, for those users who wish to generate new sequencing data sets with specific features, identification and generation of ground truth data sets is a laborious and cost-prohibitive endeavour. Moreover, it is extremely difficult to empirically determine the analytical consequences of different sample processing methods, experimental variability in library preparation and issues of sequencing bias in analysis [14].

Simulating NGS data provides an inexpensive alternative for assessing new algorithms in the context of sequencing data variation as noted [15]. With simulated datasets, one can start refining analysis procedures *in silico*. Simulated NGS datasets can incorporate the variability associated with NGS sequence data including: sequencing coverage; number of libraries and insert size; base error rates; tool parameters at the data analysis level. For in silico NGS data, a large number of SV characteristics can be readily designed including the number, the category, the size, the breakpoint sequence, the variant fraction and the haplotype for any given locus. As a result, investigators can use this simulated data to assess the potential performance and make the trade-off between analysis cost and sensitivity before even carrying out the experiment.

A number of programs generate NGS read sets to simulate metagenomics or single nucleotide polymorphisms are available [16–22]. Only recently have we seen the development and release of structural variant simulators. An early example is RSVSim [23], an R package which amends template sequence files with structural variant changes. However, it requires an interactive R session thus does not support batch processing. SCNVSim [24] improves upon RSVSim by providing a command line interface. It simulates somatic copy number variants given a number of desired SV events and/or contigs. Nonetheless, both SCNVSim and RSVSim can only output mutated contig files (FASTA), which require external steps to simulate sequence reads (FASTQ) and output resulting alignments (BAM). VarSim [14] improves upon RSVSim and SCNVSim with integrated read simulation using read simulators such as ART [25]. Instead of using a template sequence file, BAMSurgeon [26] patches an existing alignment file to embed structural variants. However, this application requires a high depth of coverage in the existing BAM file to successfully assemble a local contig for sequence patching. Moreover, the resulting structural variant may not have the exact breakpoints for the intended simulation. Overall, none of the listed tools provide a straightforward, joint control of an individual variant, including its exact breakpoints, ploidy and locus-specific allelic fraction. These features are particularly useful in simulating the clonal expansion of somatic structural variants as seen in tumors.

As a solution to the limitations of current structural variant simulators, we designed and implemented SVEngine, a full-featured simulation program suite. SVEngine is capable of generating short sequence read sets, such as produced by an Illumina system, for thousands of spike-in variants that cover different types, sizes, haplotypes and allelic fractions. Our application produces these simulated NGS data sets in a fraction of the time of other similar tools. SVEngine’s flexibility for accepting different formats enables a user to generate whole genome or targeted sequencing data mimicking germ-line, somatic and complex clonal structured genomes with ease. It offers a high degree of allelic control through its parallelized divide-and-conquer planning scheme. In the simplest mode, users only need to provide the template (reference) sequences and a desired meta-distribution of type, size and variant frequency to receive a full set of resulting FASTA, FASTQ and BAM outputs along with the ground truth BED file.

### SVEngine features and simulation performance

We compare the available features of SVEngine with other simulators that include RSVsim [23], SCNVsim [24], VarSim [14] and BAMsurgeon [26], as shown in **Table 1**. SVEngine and the other tools can simulate common types of copy number events, *e.g*. deletions and tandem duplications. All simulators except SCNVsim can simulate copy number neutral events, including insertions, inversions, and translocations. SVEngine improves the simulation of more complex SV events – it incorporates a variety of additional structural variant types originating from a combination of changes, such as inverted translocations, inverted duplications, duplicated translocation, and foreign sequence insertions. Users directly specify these events while preparing their input parameters – this process is more streamlined compared to other tools. For example, viral genome sequence insertion, which is a hallmark of the genomes of infected cells as seen in viral diseases and cancers [27], is easily achieved with SVEngine, but not available with other simulation software.

**Table 1.** Available features of structural variant simulators.

| Use Cases | SVEngine | RSVsim | SCNVsim | VarSim | BAMsurgeon |
| --- | --- | --- | --- | --- | --- |
| **Copy number events:** deletions, tandem duplications | ✓ | ✓ | ✓ | ✓ | ✓ |
| **Copy number neutral events:** inversions, insertions, translocations | ✓ | ✓ | ✗ | ✓ | ✓ |
| **Phylogenetic clonal structure:** cancer evolution | ✓ | ✗ | ✓ | ✗ | ✗ |
| **Foreign Sequence Insertion:** virus integration | ✓ | ✗ | ✗ | ✗ | ✓ |
| **Non-human genome:** various template and ploidy | ✓ | ✓ | ✓ | ✗ | ✓ |
| **Not requiring pre-existing alignment:** no high coverage BAM input | ✓ | ✓ | ✓ | ✓ | ✗ |
| **Generate mutated contig:** FASTA output | ✓ | ✓ | ✓ | ✓ | ✗ |
| **Generate short reads:** FASTQ output | ✓ | ✗ | ✗ | ✓ | ✗ |
| **Generate alignment:** BAM output | ✓ | ✗ | ✗ | ✗ | ✓ |
| **Locus-specific variant ploidy** | ✓ | ✗ | ✗ | ✗ | ✗ |
| **Locus-specific variant frequency** | ✓ | ✗ | ✗ | ✗ | ✓ |
| **Exact breakpoint** | ✓ | ✓ | ✗ | ✗ | ✓ |
| **Multiple sequencing libraries** | ✓ | ✗ | ✗ | ✗ | ✗ |

In terms of input/output flexibility and ease of use, SVEngine provides automates template sequence modification, read simulation and read mapping steps. These features are not found in other simulators of SV events. Also, SVEngine is the only tool which outputs a full set of simulation results in standard formats, including altered contig sequence (FASTA), simulated short reads (FASTQ) and alignment (BAM) files **(Figure 1)**. At the input step, all tools take in template sequences in FASTA format as the starting material, while BAMsurgeon additionally requires a pre-existing alignment file in BAM format as input. Overall, read coverage of this BAM file has to be large (typically >30x), to successfully assemble local contigs. Such requirements preclude the use of BAMSurgeon in applications generating low coverage and consequently limit its users to mimicking conditions based on available high-coverage BAMs. The VarSim tool needs structural variant prototypes from DGV [28] making it only applicable to the human genome. At the output step, RSVsim and SCNVSim provide modified sequence contigs in FASTA files. BAMsurgeon outputs modified alignment in BAM files, but without providing associated modified contigs. VarSim provides both contigs containing a variant and simulated short reads, but it still requires additional user effort to generate alignment files.

**Figure 1.**
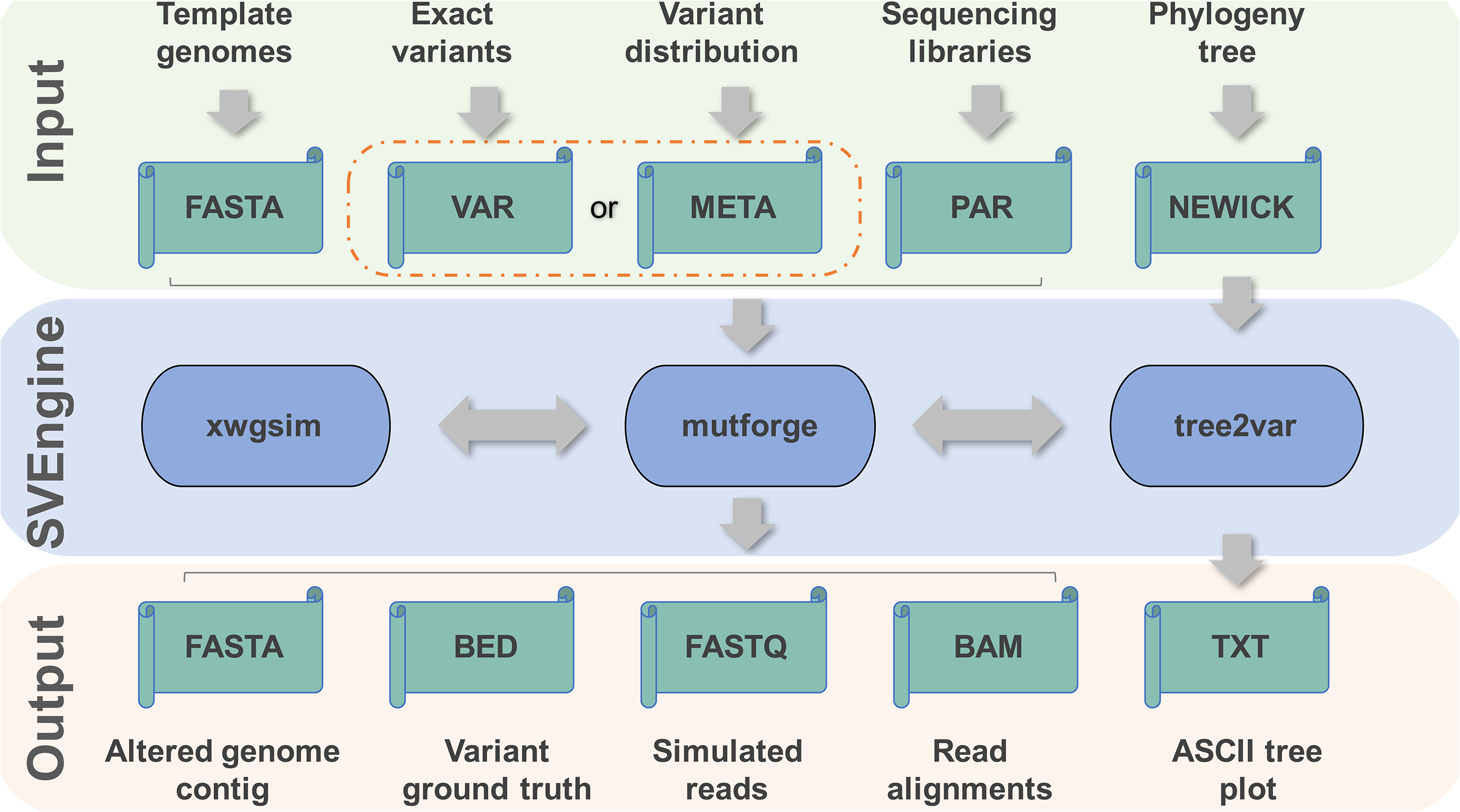
Inputs, outputs and execution components of SVEngine. The flow of data was marked by gray arrows. The input, SVEngine functioning and output data spaces were color shaded.

With regard to precise and versatile control of individual variants, SVEngine enables one to easily specify variant type, size, exact breakpoint, ploidy and allelic fraction for individual loci. Additionally, SVEngine simulates a full spectrum of germline, somatic and clonal structural variations by the specified meta-distribution. In comparison, RSVSim does not support loci-level control, as it only patches template sequence on demand. With SCNVsim and VarSim, one can only control a meta-distribution of structural variants, such as the total number for each variant type, minimum and maximum variant size. SCNVsim allows the specification of ploidy, number and type of clones but does not have the capability to specify exact breakpoints. VarSim randomly resamples breakpoint and other variant information from a DGV database dump. Only BAMsurgeon and SVEngine support locus-specific variant fractions, *i.e.* allowing different allele fractions for individual variants. Only SVEngine supports locus-specific ploidy, *i.e.* allowing a different ploidy state for individual variants. Both BAMsurgeon and SVEngine also support exact breakpoints for individual variants. However, in practice, the actual breakpoints generated by BAMsurgeon may differ from input, as a result of improvised local contig assembly. Another unique feature of SVEngine is the ability to specify multiple sequencing libraries, which can each have different insert size mean and standard deviation, intended coverage depth, and read length.

In addition to the features listed in **Table 1**, SVEngine allows user to designate some regions while avoiding others. Examples of such applications include simulating exome or targeted sequencing data sets. Moreover, this feature enables one to avoid complex regions such as telomeres and centromeres. SVEngine also features parallelized simulation by dividing genome into pieces, embedding variants into each piece and then stitching them together. Therefore, its performance can be boosted using the multi-core processors.

**Table 2** lists the runtime on a test set of 15,000 SV events into a 30x coverage whole genome sequencing simulation, including 2,500 events each of deletion, tandem duplication, inversion, translocation and domestic or foreign sequence insertion. In multi-processor mode, SVEngine has the shortest runtime in all three levels of simulation, i.e. obtaining altered contigs, simulated reads and alignments in FASTA, FASTQ and BAM formats in less than 10 min, 20 mins and 3 hours, respectively. Overall, SVEngine is 1x, 15x and 48x times faster than the single-process SVEngine run. The performance scales almost linearly with the added CPU power in generating the alignment output, because the read mapping time cost dominates other time costs including data serializing time. Moreover, even the single-process SVEngine (SVE-single) is more efficient than its other counterparts. For example, it took only 10 mins to SVE-single to generate all altered contigs when RSVSim and SCNVSim took several hours. SVE-single required half the time BAMSurgeon needs to generate all read alignments. All run time were all measured on 8 Intel^®^ Core i7-4790 CPU @ 3.60GHz with 32 GB RAM computer.

**Table 2.** Runtime performance comparison.

| 15000<br>events at<br>30x<br>coverage | SVEngine<br>(64 procs) | SVEngine<br>(1 proc) | *RSVSim* | *SCNVSim* | *VarSim* | *BAMSurgeon* |
| --- | --- | --- | --- | --- | --- | --- |
| FASTA<br>output | <10 min | < 10 min | 10 hr | 2 hr | Not<br>Available | Not Available |
| FASTQ<br>output | <20 min | 5 hr | External | External | 6 hr | Not Available |
| BAM output | 2hr | 5 days | External | External | External | >10 days |

**Not Available:** output format is not available
**External:** output format is only available through additional external software, thus not tested

### Simulating cancer genome evolution

SVEngine provides a high degree of control over SV events with variable allelic fractions – this feature enables one to simulate heterogeneous cancer genomes undergoing a phylogeny tree-structured clonal evolutionary process. As a demonstration, we present a simple example scenario in which we simulated with SVEngine (**Figure 2**). To simplify the description of the phylogenetic process of cancer evolution, we use a binary tree representation of phylogeny. This binary tree is easily converted to a typical phylogeny tree by merging all nodes of identical cell subpopulations.

**Figure 2.**
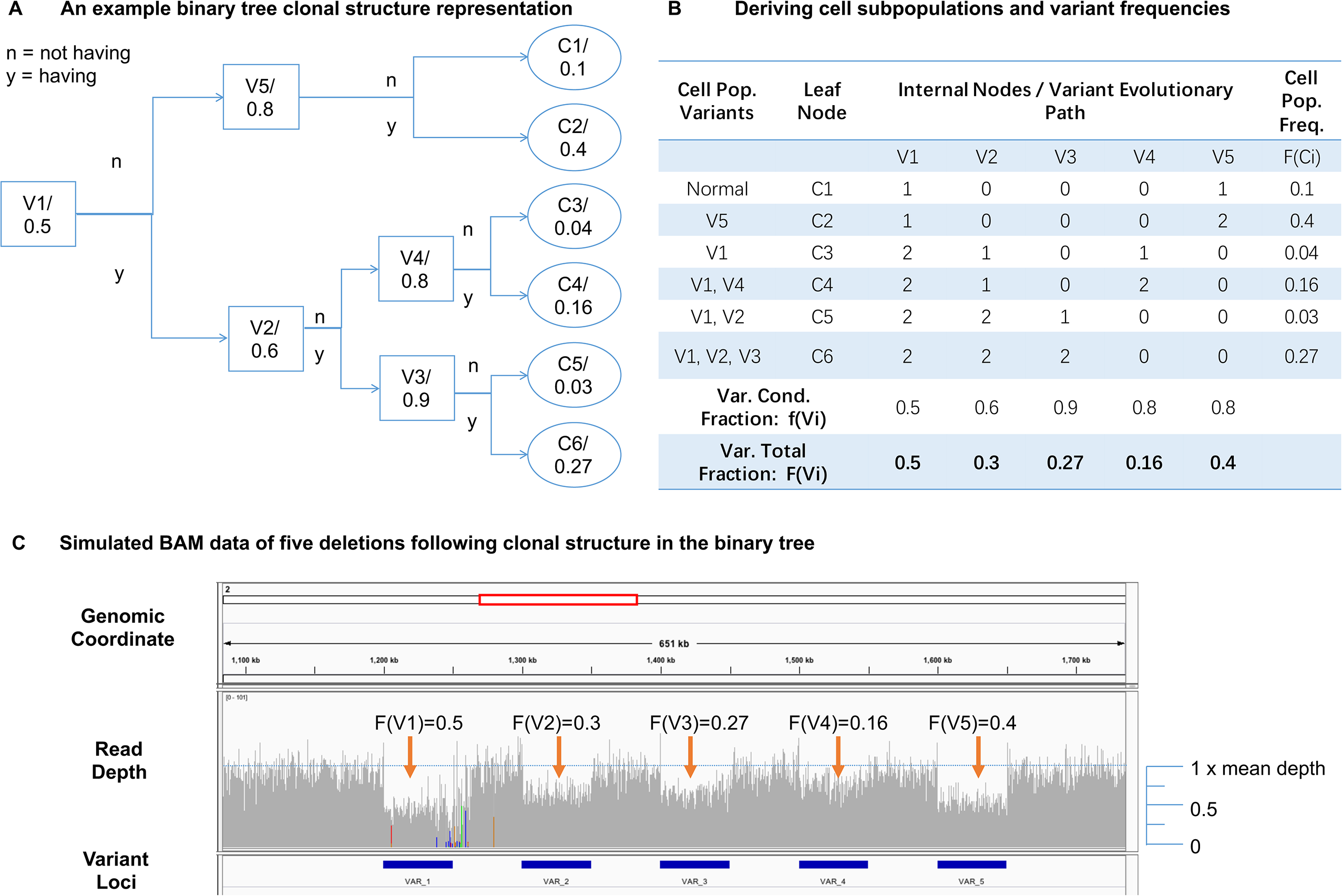
Simulating cancer evolution. **(A)** An example cancer evolution tree. The conditional fraction in each internal node represents the fraction of cell population gaining the next structural variation, which is represented by the label of the internal node. **(B)** An example computation table to determine final variant frequency of each variation and cell population frequency of each terminal genotypes. **(C)** Integrated Genomics Viewer view of SVEngine simulated BAM data of five deletions following clonal structure in the example binary tree.

One example is a binary tree shown in **Figure 2A**, where each of the five (*m* = 5)internal tree nodes denotes a bifurcation event when part of the parental cell population is gaining an additional mutation (*V_j_,j = 1…m*) The root node represents the lowest common ancestor genotype of all subpopulations of cancer cells. It is typically the normal germline cell as depicted here, or the first generation of cells bearing common somatic mutations that start becoming cancerous. The “root” cell populations are split by the next immediate event, i.e., gaining the mutation *V*_1_, resulting in two child populations depending on a cell’s status of carrying *V*_1_ or not, as represented by its two child nodes. We denote the conditional cell fraction of gaining *V*_1_ as *f(V_*1*_)*, which is 50% or 0.5 in this case, and is denoted at the root. The mutational process goes on for subsequent internal nodes and until all variants (a total of five in this example) are represented by their bifurcation internal node. The resulting binary tree has six *(n = 6)* leaf nodes (*C_*i*_,i = 1…n*), which represent all possible somatic genotypes of the terminal cell subpopulations.

As we can see, any terminal somatic genotype is completely determined by following the mutational path from the root down to a leaf node. We use a tertiary vector *C_*i*_= (C_*i,1*_…C_*i,m*_*), to indicate such path, where

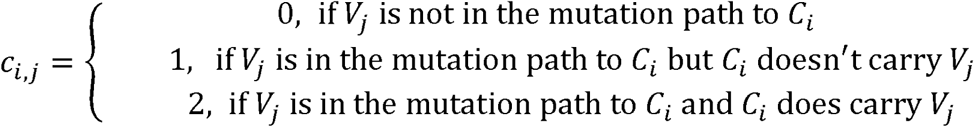

In addition, we define the conditional frequencies *f(V_i_)*, which is the fraction of cells derived from a parent population carrying event *V*_*i*_ as: 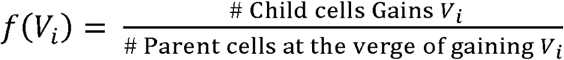 Therefore, the final population frequency *F(C*_*i*_) of cell subpopulations *C_i_* is expressed as:

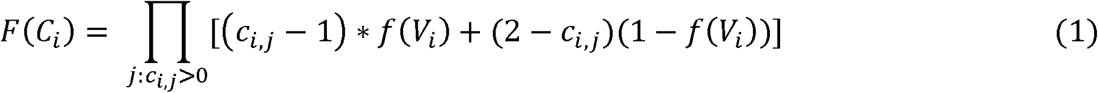

With 1(.)as the indicator function, the concurrent proportion *F*(*V_j_*) of all extant cells is simply the marginal sum of all cells carrying *V_j_*:

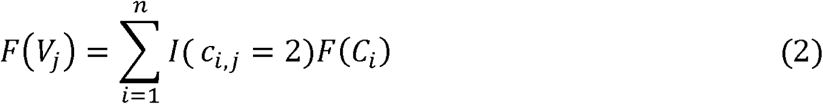

**Figure 2B** shows the derivation of the above quantities for the example binary tree. The sequence of events ensures a partial order that the mutant allele frequency is always higher for events occurring upstream, as compared to events occurring downstream on the same lineage. It is possible that terminal genotypes may not all coexist in extant populations. The proposed binary tree representation accommodates a deceased population by having zero proportion for such leaf node. SVEngine allows user to input a binary tree with relevant bifurcation fractions to structure the variant fractions that falls along the line of the evolutionary tree. For designating this feature, the input to SVEngine is in standard NEWICK format – a widely accepted format using parenthesis to encode nested tree structures [29]. Each internal node is labelled by the population splitting variant and weighted by the conditional splitting fraction. Each leaf node is labelled by associated terminal genotype and weighted by the subpopulation fraction as an optional feature. For instance, the NEWICK string for the example binary tree is: ((C1, C2) V5: 0.8, ((C3, C4) V4:0.8, (C5, C6) V3: 0.9) V2:0.6) V1:0.5.

**Figure 2C** shows the IGV browser view of SVEngine simulated BAM alignments of five equal-size deletions following the mutational process as represented by the example binary tree. The read depth shows the difference of allelic fractions corresponding to the computed final variant fractions based on the tree. We display an example of monoclonal cancer evolution, assuming that all cell subpopulations start from a set of common ancestor cells (as denoted by root node in the tree). However, simulations of multiclonal evolution are also possible with SVEngine. For example, one simply assigns an empty event to *V*_*1*_ and then sets the conditional fractions of the two child events *V*_*1*_ and *V*_*5*_ to 100% to simulate a two-clonal origin evolution. With SVEngine’s high efficiency, the simulation is easily scaled to tens of thousands of variants with a tree having a more complex structure.

### Simulating the multitude of structural variations

Current structural variation detection methods mostly rely on detecting altered read mapping features to identify structure changes [30]. The most important such features are read depth/coverage, read pair insert size, single ended read pairs (hanging reads), soft-clipped reads and split reads (clip/split reads). It is essential for structural variant simulators to correctly produce such feature changes corresponding to the causal event. In **Figure 3**, we comprehensively illustrate the expected changes in mapping that result from different types of structural variants.

**Figure 3.**
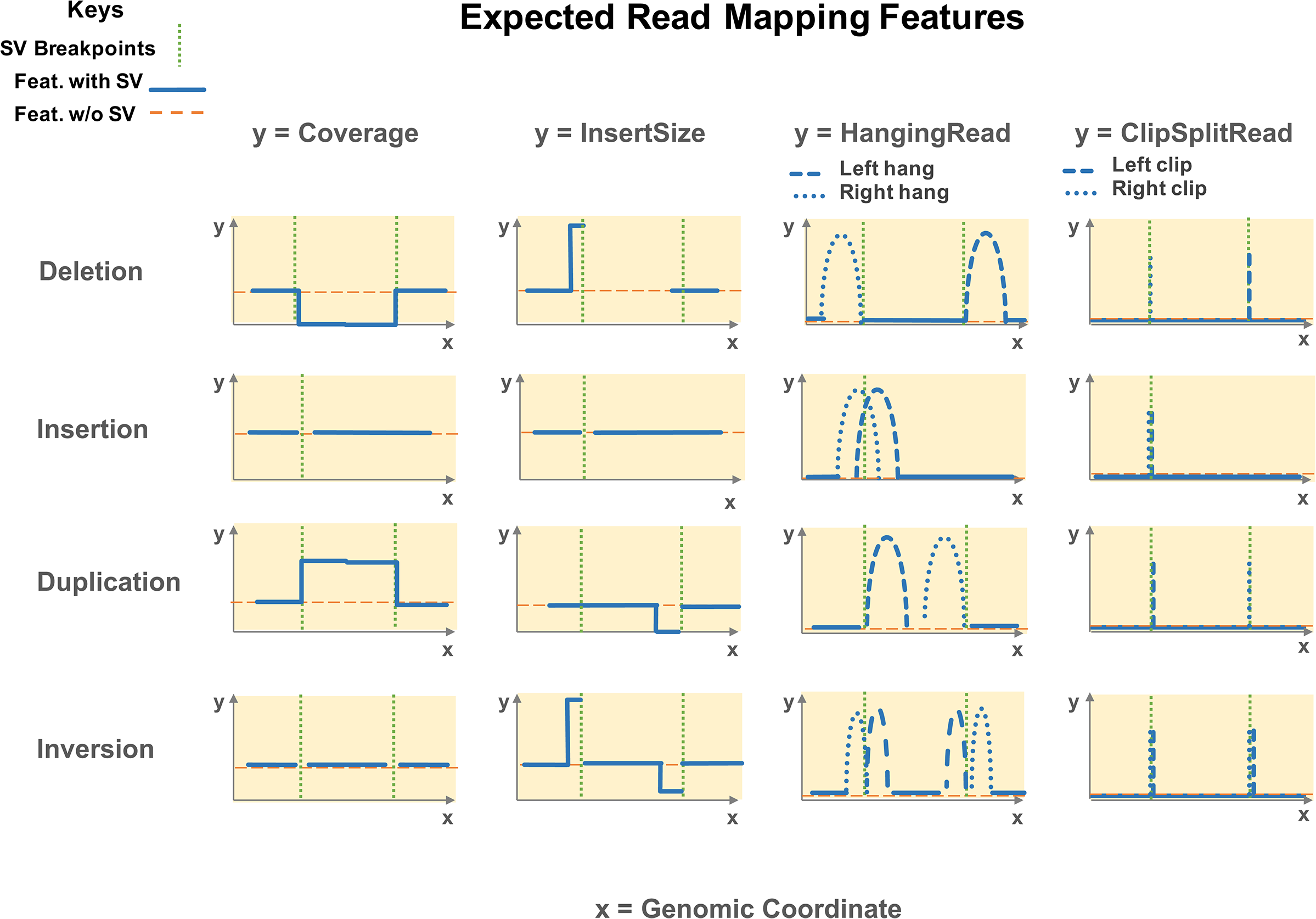
Expected read mapping features of structural variant prototypes. Rows – variant prototypes: 1) Deletion, 2) Insertion, 3) Duplication, 4) Inversion. Columns – mapping features: 1) Read coverage, 2) Read pair insert size, 3) Single end mapped read (HangingRead), 4) Soft clipped read or split mapped read (ClipSplitRead). X-axis is genomic coordinates. Y-axis is feature value/counts. Dashed orange line stands for expected feature value without alteration. Solid blue line stands for expected feature value with alteration. Dotted green bar denotes the breakpoint(s).

In the scenario of a deletion (**Figure 3**, **first row**), all the mapping features, such as coverage, insert size, hanging read and soft-clip/split read are expected to change, as illustrated in the *Coverage*, *InsertSize*, *HangingRead* and *ClipSplitRead* columns, respectively. First, there is a reduction of read coverage over the deleted region because no reads are present. Second, the insert size of read pairs that are mapped straddling the breakpoints is expected to increase as inferred by alignment to the reference. This extended insert size is possible because the deleted region is not present in the real DNA molecules where these read pairs originating from. Third, a fraction of read pairs aligning to the left of the left breakpoint lack a mapped mate read – this forms right mate hanging read pairs (right hang). It is because the left breakpoint has interrupted the mate mapping by reducing similarity between the read and the reference. Due to symmetry, left mate hanging read pairs (left hang) form to the right of the deletion. Finally, when the breakpoint interruption in the mate is not as severe, it is possible that the mate read can still partially map. The noncontiguous part of the mate is either clipped or, if it is long enough, mapped to near the other end of the deletion. Such resulting read pairs are what we refer to as left or right soft-clipped (or split mapped) reads depending on which side of the reads were split or clipped. These read pairs are expected to map right next to both breakpoints with the clipping (or splitting site) aligned to the exact break point location, as shown.

For an insertion (**Figure 3**, **second row**), the most noticeable change is the clustering of both the right and left hanging read pairs centering over the breakpoint. One observes a similar clustering for the left and right clip/split reads. As shown in **Figure 3**, an insertion exhibits fewer changes than other types of structural variants, and so insertions are generally the most difficult to detect. In the scenario of a tandem duplication (**Figure 3**, **third row**), the read coverage is expected to increase within the duplicated region. The insert size of reads mapped to the left of the right breakpoint is expected to decrease, or even be negative, because the mate is likely to have the same sequence as the segment preceding the read. Then, when the mate is mapped upstream of the current read, it causes a reversal of normal read strand order and introduces a negative insert size in the read mapping. By the same reasoning, the right hanging, clipped and split reads are clustered upstream next to the right breakpoint. Similarly, the left hanging, clipped and split reads are clustered downstream next to the left breakpoint, making the tandem duplication almost a mirror image of deletion.

In the scenario of an inversion (**Figure 3**, **fourth row**), the coverage shows almost no change. The insert size near the left breakpoint is similar to the deletion scenario, which has an increase. This is because the mate is from the reverse complement of the other end of the inverted segment, which creates an inflated insert estimate and an abnormal forward-forward strand read pair. Similarly, the insert size near the right breakpoint is decreased and forms an abnormal reverse-reverse strand read pair. When these abnormal pairs are interrupted by the breakpoints, it creates corresponding hanging read and clipped/split read clusters around both breakpoints. Citing another example, a chromosomal translocation is simply a combination of features at the region deleted by the translocation and insertion features at the region inserted.

## CONCLUSION

We have developed and released SVEngine, a structural variant simulator, available as an
open source program. It simulates next generation sequencing data that has embedded structural variations as well as an assortment of complex sequence features. SVEngine simulates and outputs mutated sequence contigs (FASTA), sequence reads (FASTQ) and/or alignments (BAM) files with desired variants, along with BED files containing ground truth. SVEngine’s flexible design enables one to specify size, position, and heterogeneity for deletion, insertion, duplication and inversion and translocation variants. SVEngine’s additional features include simulating sequencing libraries having multiple different molecular parameters, and targeted sequencing data sets. SVEngine is highly parallelized for rapid and high throughput execution.

We showed the versatility and efficiency of SVEngine by comparison of features and runtime versus other available simulators. We demonstrated the utility of SVEngine in an example mimicking the phylogeny in cancer clonal evolution, by simulating the associated variant allelic frequency. We validated the accuracy of SVEngine simulations by examining expected sequence mapping features such as coverage change, read clipping, insert size shift and neighbouring hanging read pairs for representative variant types. SVEngine is implemented as a standard Python package and is freely available for academic use at: https://bitbucket.org/charade/svengine.

## METHODS

### Simulation software and pipeline

SVEngine was developed as a standard Python package with a C extension. SVEngine provides two Python executables and one C command line executable: *mutforge*, *tree2var* and *xwgsim*, respectively. The *mutforge* command implements a parallelized algorithm that divides the template genome into blocks of contigs, spikes structural variants into the contigs, samples short reads from the altered contigs, and finally merges the short-read sets back into one file and performs the alignment. The *tree2var* command implements a procedure that determines variant fractions from an input phylogeny tree based on **Equations (1)** and **(2)** and a depth first search graph algorithm, and then substitutes these allele fractions in an input VAR file. The *xwgsim* command implements a modification to *wgsim*, which reduces the read sampling rate by 50% for the overlapping regions between contigs (i.e. ligation regions). The overlaps were designed so as to allow for the proper merging of contig-wise read sets. *xwgsim* only interacts with *mutforge* thus is mostly transparent to a user.

As shown in **Figure 1**, the required inputs to *mutforge* are three-fold: 1) a template haploid sequence file(s) in FASTA format. This can be a standard human genome reference, or any other reference genome sequence. 2) A VAR file or a META file for specifying structural variants (distributions). These are tab delimited files with columns defined in SVEngine’s manual. The VAR format is intended for specifying exact information for individual variants, which includes variant id, parent id (if part of a complex event such a deletion occurring due to a translocation), fraction, ploidy, chromosome, starting position, and the sequence length to be deleted and/or the sequence content to be inserted. Alternatively, the META format is intended for higher-level control, allowing one to specify a desired meta distribution of variants, including variant type, and total number of events, size, allele fraction, and ploidy distributions per type.

One can specify where and how to insert the sample sequence in the case of foreign DNA insertion. For example, a user can readily design 100 deletions of size ranging from 100bp to 10kbp, of a uniform distribution of allelic fraction and a fair Bernoulli distribution of homo– and heterozygosity in one line of text in the META file. 3) The PAR file is used to model an experimental design, including insert size, read length and coverage, as well as additional options for *xwgsim*. The file can be used to specify multiple libraries with different mean insert size and standard deviation. One can use such normal mixtures to approximate irregular libraries of multiple modes and asymmetric tails. The *xwgsim* command also provides random embedding of SNVs and indels if desired. In the SVEngine’s Wiki page, we supply example VAR, META and PAR files with detailed annotation to facilitate their usage.

Once all inputs are provided, the SVEngine master process divides the template genome into blocks and serializes spike-in tasks to parallel worker processes. The worker process patches its assigned contig and if read pairs were required, it also calls *xwgsim* to simulate read pairs. The read pair subsets are then collected by the master process and merged, and if alignments were required, it also calls *bwa-mem* and *samtools* to map the reads to the reference.

The output of SVEngine has three levels. At the first level (contig), only two files would be generated: one is a FASTA file containing all the altered contigs, the other is the ground truth of spiked-in variants in a BED3 format file with the additional columns following the VAR format as in the input. At the second level (read pair), SVEngine additionally outputs the read 1 and read 2 of the simulated read pairs in two FASTQ files. Finally, at the third level (alignment), SVEngine provides the read alignment output to the given reference in a BAM file format. The runtime of SVEngine increases with the specified output level, as additional processing time will be required. Table 2 can be used as a reference for runtime estimates for different output levels.

The *tree2var* command simulates a clonal evolution scenario, which requires an additional tree input file (NEWICK). *tree2var* also takes a VAR file, which can be generated from *mutforge* with a META file input and in the dryrun mode. The user must insure that the identifier of the tree’s internal nodes and the variant match each other, as this is used to identify and replace allele fraction with the value computed from tree phylogeny. The *tree2var* outputs a new VAR file which contains the rewritten allele fraction fields that reflects the clonal structure described by the user tree. For intuitive diagnostics, *tree2var* also outputs a ASCII text based plot of the parsed input tree. SVEngine’s tree parsing interacts with DendroPy [29], which allows further functionality such as random tree simulations and many tree statistics. The output VAR file from *tree2var* then becomes the input to *mutforge* for actual read simulation.

### A parallel simulation framework

SVEngine’s major improvements to existing structural variant simulation tools involve one’s ability to alter the allelic fraction, control of haplotypes and highly efficient parallelized simulation. These improvements were achieved through the core algorithm as illustrated in **Figure 4**. In general, we used a divide-and-conquer approach intertwined with multi-process execution: First, the SVEngine master process lays out a genome grid for simulation. For any input haploid sequence, the entire genome is partitioned into *N* equal size non-overlapping blocks: *B_1_,B_2_,…,B_N_,* where 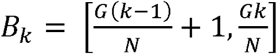. The block size 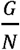 can be chosen at the input, where *G* is the entire genome length. Ligation regions of length *l* are also defined, which consists of symmetric touching border regions of equal size adjacent blocks: *L*_*1*_*,L*_*2*_*,…,L*_*N-1*_
, where 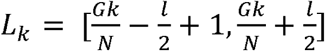. These serve as buffer regions that enable the SVEngine to ligate block sequence-based simulations back together. The block generating procedure is similar for multi-chromosome genomes except blocks representing chromosome ends might be shorter than the standard block size.

Second, the SVEngine master process coordinates all of the tasks. In one task, a structural variant is embedded into the adjacent sequence – this is done by assigning a sequence of blocks that it impacts. All the variant’s control information is attached to the task as well. In the figure, the first variant, a 50% deletion *SV*_*i*_was assigned blocks 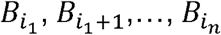 and the next variant, a 100% deletion *SV*_*j*_ was assigned blocks 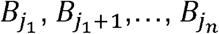. Depending on its size, a variant can take anywhere from one block, to as many blocks as needed. The genomic region which is not altered, between adjacent variants, say *SV*_*i*_ and *SV*_*j*_, also becomes a task. This is assigned to the sequence blocks complementary to the blocks taken by *SV*_*i*_ and *SV*_*j*_ and with a no-op instruction attached. If necessary, no-op tasks with large block sequences are further broken down to no-op tasks with size-capped block sequences to improve efficiency of parallelization.

**Figure 4.**
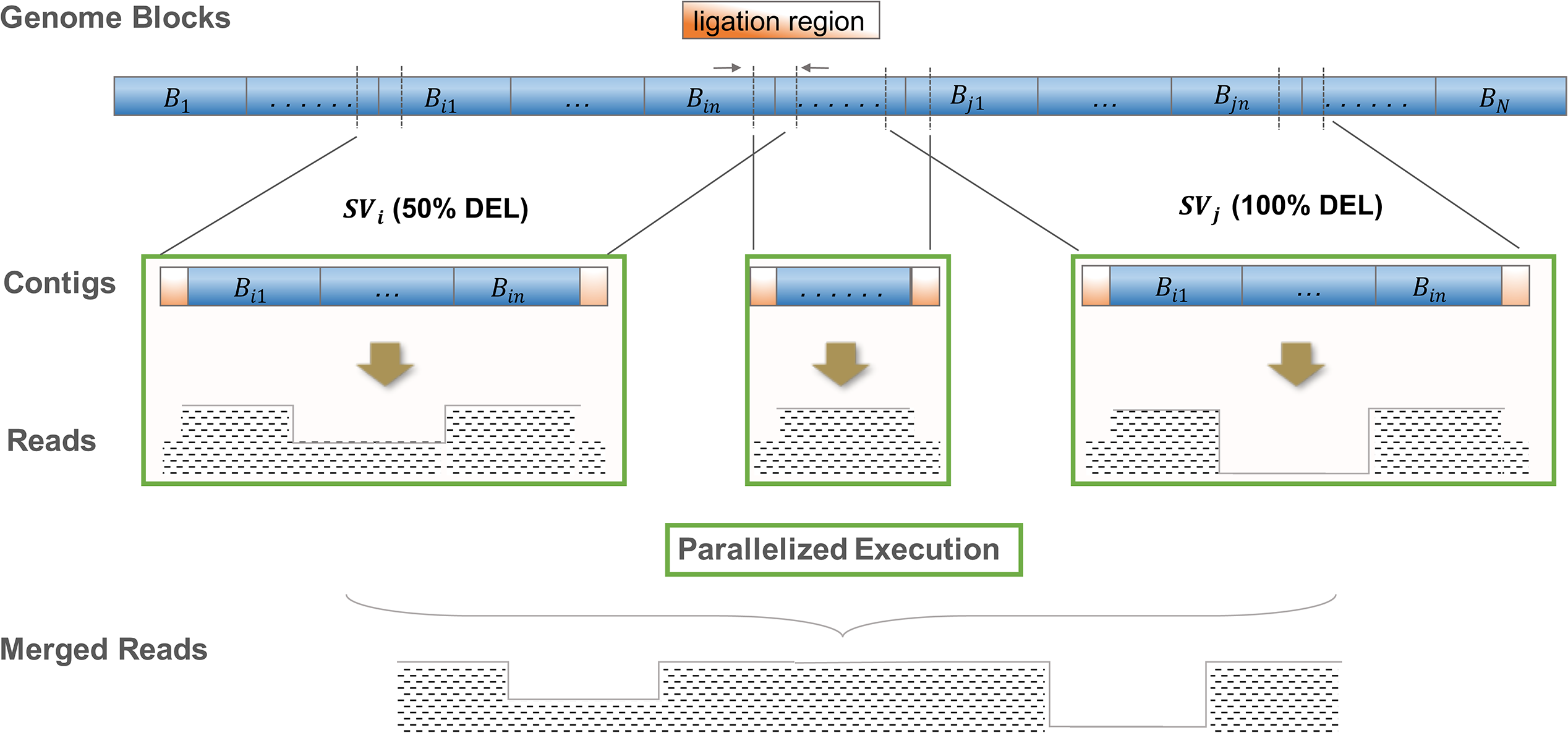
The core parallelized simulation algorithm of SVEngine. Here are two neighboring events 
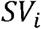
and 
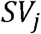
a 50% deletion and a 100% deletion to be spiked-in. The first deletion event spans blocks 
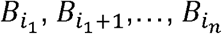
 and the second deletion event spans genome blocks 
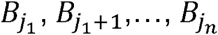
 The genome blocks are shaded in blue while the ligation regions are shaded in orange. The resulting read pairs are represented by their coverage in black dash patterns. The parallel execution tasks were boxed in green color.

Third, the SVEngine master process dispatches all the tasks to an auto revolving worker process pool and then waits for all the workers to finish. Each worker process, when assigned a new task, loads the haploid sequence defined by the task’s block sequence plus left and right ligation regions. For example, a worker would load sequence from 
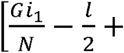

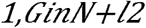
 for *SVi* as the original contig, or *Gj1N−l2+1, GjnN+l2* for *SVj*, or 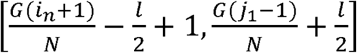 for the no-op task in between them. The original contig is then operated on for deletion, insertion, or other alternations to form the altered contig. If no-op the original contig is unaffected. The worker then calls *xwgsim* to simulate the proper numbers of read pairs from the original and altered contigs according to the specified frequency and resulting contig sizes. The *xwgsim* step also takes care of attenuated sampling (at half the normal rate) within the designated ligation regions as the worker provides the ligation size *l* in its arguments. In addition, *xwgsim* adds a procedure to the popular NGS simulator *wgsim* [31], which rejects a new read pair at 50% chance if any of its two ends originates in a ligation region. Patterns of expected read pair coverage from the *SV*_*i*_, *SV*_*j*_ and no-op tasks are illustrated in **Figure 4**.

Fourth, when the worker processes are completed, the master process collects all simulated read pairs from all tasks and concatenates them into two final files, one for read 1 and the other for read 2. Also, it collects all original and altered contigs and concatenate them into one final sequence file. Finally, it performs read pair alignment to the reference genome using *bwa-mem* and samtools. This is last step, although sequential in SVEngine, is already thread parallelized by other required programs such as the *bwa* and *samtools* tools [22, 32]. Patterns of expected read pair coverages after merging the *SV*_*i*_, *SV*_*j*_ and no-op tasks are also illustrated in **Figure 4**. The described algorithm assumes one haploid for simplicity. For multi-ploidy, each haploid is handled in a similar way by the worker process except that the variant’s haplotype status is also taken into consideration. Overall, this SVEngine’s core algorithm is very efficient as demonstrated by the runtime comparison, is very versatile and isaccurate as demonstrated by multiple example applications described in this paper.

### Notable simulator features

To comprehensively evaluate structural variant callers, one may need a wide spectrum and large number of SV events. This range is more easily specified by distributions of variants rather than individual variants. SVEngine supports variant distributions as specified in the META format. The expansion of distributions to actual variants takes place in the master process before any spike-in. The distributions are expanded on a target genome sequentially by randomly pick the next event’s start position from regions that can accommodate it. Afterwards, it removes the impact region from the remaining available regions and so on. Once all distributions are expanded, the master process returns a list of variant fulfils user’s specification and outputs them into a VAR file. The user can choose to run SVEngine in dryrun mode to stop the execution at this point and inspect the resulted variants. The user also has the option to continue the simulation to the end, which is equivalent to input the output VAR file into SVEngine for simulation in the next step.

To increase the sensitivity of SV detection, researchers may prepare multiple sequencing libraries with different molecular parameters for analysis. For example, different insert sizes enable the detection of a wider spectrum detection of SVs [33]. Longer sequence read length can boost the performance of some callers that employ remapping strategies [34]. A unique feature of SVEngine is its ability to simulate NGS data modelling multiple libraries with different mean insert size and standard deviation, coverage and read lengths. The feature is implemented within the worker process. When using a multi library task, SVEngine will call *xwgsim* multiple times to generate read pairs in accordance with the library specification.

SVEngine provides simulation data that target or masks specific genomic regions. This feature emulates targeted sequencing applications, such as exome sequencing and gene panel sequence data. It can be used to exclude problematic regions such as gaps, telomere and centromere regions of the reference template. One only needs to provide standard BED format files to SVEngine listing the regions to be masked or targeted by the simulated sequencing.

Databases like DGV and much other literature provide a list of known population variants. Tools like RepeatMasker (http://repeatmasker.org) provide extensive lists of known regions of human repeats and/or homolog sequences, with enrichment of structural variant breakpoints. Although not provided in our examples due to their varied formats, in principle, these population and repeats-mediated variants can be downloaded in general tab delimited formats, such as BED or VCF files. Subsequently, these annotation formats are easily converted into an SVEngine VAR format input file using text processing utilities such as *awk* and *sed*. Using a VAR file generated in this way, SVEngine can easily embed these variants into simulation data.

## Availability of supporting source code and requirements

Project name: SVEngine

Project home page: https://bitbucket.org/charade/svengine

Operating system: Linux/Unix

Programming language: standard Python package with a C extension

Other requirements: GNU C Compiler or similar

License: Stanford University

## Data availability

In silico data sets are available at https://bitbucket.org/charade/svengine.

## Abbreviations

SV: Structural Variation/Variant
NGS: Next Generation Sequencing

## Ethics approved and consent to participate

Not applicable. No human subjects or data. No animal subjects or data.

## Consent for publication

Not applicable.

## Author contributions

LCX, NRZ and HPJ designed the study. LCX developed the algorithm, wrote the program and tested the software. HJL and NA provided feedback on the software design and testing. LCX and DMA generated and analyzed the data with assistance from CL. LCX and HPJ wrote the manuscript. All authors approved the manuscript.

## Acknowledgements

We thank John Bell, Sue Grimes and Erik Hopmans at Stanford University and Yuchao Jiang at University of Pennsylvania for helpful discussions.

